# Zygosity-Aware DNA Language Modeling Improves Ancestry and Gene Expression Prediction

**DOI:** 10.1101/2025.11.19.689326

**Authors:** Hussin El Rashidy, Ali Saadat, Jacques Fellay

## Abstract

DNA language models (DNA-LMs) are transforming how genomic sequence information is represented and interpreted. Yet most current approaches treat DNA as a single sequence, overlooking the diploid structure and zygosity information that distinguish the two parental copies of the genome. Here, we systematically evaluate explicit diploid, zygosity-aware representations in DNA-LMs for two downstream tasks: ancestry classification and gene expression prediction. For ancestry, we use HyenaDNA embeddings of the extended MHC region and show that concatenating maternal and paternal haplotype embeddings consistently improves predictive performance across five superpopulations compared to single-haplotype inputs. For gene expression, we compare convolutional neural networks (CNNs) trained from scratch with Nucleotide Transformer models using reference-only, single-copy, and two-copy (zygosity-aware) sequence encodings. CNNs showed increased performance by incorporating genetic variation and zygosity via simple additive genotype encoding, whereas naïvely injecting variation into pretrained Nucleotide Transformer models yields mixed effects, highlighting a mismatch between current pretraining objectives and variation-sensitive prediction. Together, our results demonstrate that zygosity-aware representations can capture biologically meaningful information beyond reference-only views and underscore the need for diploid- and population-aware pretraining strategies in future DNA-LMs for variant interpretation and precision medicine.

## 1 Introduction

Advances in DNA language models (DNA-LMs) have begun to transform how genomic sequences are represented and interpreted (Benegas et al., 2025; Sanabria et al., 2024). By adapting architectures originally developed for natural language processing, such as Transformers and related sequence models, these methods learn contextual representations of DNA that can be reused across diverse predictive tasks, including gene expression modeling, chromatin state prediction, and variant effect interpretation (Avsec et al., 2021; Benegas et al., 2023). Pretrained models such as the Nucleotide Transformer (NT) (Dalla-Torre et al., 2024) and HyenaDNA (Nguyen et al., 2023) are typically trained in a self-supervised fashion—using objectives like masked language modeling or next-token prediction—to capture patterns in large corpora of genomic sequence.

Most current DNA-LMs, however, adopt a fundamentally haploid view of the genome. They operate on a single sequence derived from a reference assembly or a consensus haplotype, and therefore do not explicitly model the diploid nature of human genomes. Two key sources of biological signal are thus underutilized. First, genetic variation (GV) which reflects population-level diversity, including single-nucleotide variants (SNVs) and other polymorphisms that differ across individuals and ancestries. Second, zygosity which describes whether the two alleles at a locus are identical (homozygous) or different (heterozygous), and is central to dominant and recessive inheritance patterns (Wilkie, 2018; Saadat & Fellay, 2024). Ignoring these dimensions may limit performance precisely in tasks where variation and dosage matter most.

In practice, many large-scale DNA-LMs are pretrained primarily on the human reference genome or on a small number of haplotypes. As a result, they capture rich contextual and motif-level information but do not systematically encode how alleles differ between the two parental copies of the genome. This design choice stands in contrast to the well-established role of heterozygosity and allele frequency in population genetics and complex trait mapping. For example, heterozygosity ratios differ across human superpopulations and have been proposed as robust signatures of ancestry (Auton et al., 2015; Samuels et al., 2016). Likewise, expression quantitative trait loci (eQTLs) and other regulatory variants often act in an allele-specific manner, with zygosity modulating effect size and penetrance (Lappalainen et al., 2013a; Huang et al., 2023). These observations suggest that diploid-aware sequence representations could provide additional signal beyond what is available from a single, reference-like sequence.

Ancestry prediction offers an ideal test case for diploid modeling. The MHC region on chromosome 6 is one of the most polymorphic areas of the human genome, harboring highly diverse HLA and immune-related genes that differ across populations (Auton et al., 2015; Dilthey, 2021). Patterns of alleles and heterozygosity within the MHC reflect demographic history and immune-related selection pressures. We hypothesize that explicitly modeling both haplotypes with DNA language models, rather than collapsing to a single sequence, should improve ancestry classification, particularly when using long-range encoders like HyenaDNA.

Gene expression prediction provides a complementary evaluation, focusing on regulatory effects rather than population structure. Genetic variants near transcription start sites (TSSs), notably eQTLs, can drive allele-specific expression differences. Standard models trained on the reference genome may miss important zygosity and phase information, motivating a systematic assessment of whether diploid-aware, variation-inclusive sequence representations can improve prediction of gene expression from promoter regions.

Here, we evaluate the importance of explicitly modeling both haplotypes in DNA language models for ancestry and gene expression prediction, contrasting simple genotype encodings with large pretrained models. Our study provides new evidence for the benefit of diploid-aware representations and highlights directions for future improvements in variation-sensitive genomics.

## 2 Methods

### 2.1 Data sources

#### 2.1.1 1000 Genomes Project

For ancestry prediction, we used phased variant calls from 2,548 individuals in the 1000 Genomes Project (Auton et al., 2015), focusing on the extended MHC region of chromosome 6 (28,510,120– 33,480,577, hg38) (Dilthey, 2021). Individuals were assigned to five superpopulations (AFR, EUR, AMR, SAS, EAS) based on sample metadata. Diploid sequences, including single-nucleotide variants (SNVs), were reconstructed from VCF data by extracting phased genotypes and applying variants to the reference genome. Haplotype A corresponded to the left column and haplotype B to the right column of the GT field, with variants stored in `chr:pos:ref:alt` format for reproducibility and direct use with GenomeKit. This region was selected for its high genetic diversity and relevance to ancestry due to the presence of polymorphic HLA genes (Arrieta-Bolaños et al., 2023; Buhler & Sanchez-Mazas, 2011).

#### 2.1.2 Geuvadis dataset

For gene expression prediction, we used the Geuvadis dataset (Lappalainen et al., 2013b), which contains phased whole-genome sequencing (WGS) data and log-transformed gene expression levels from lymphoblastoid cell lines (LCLs) for 455 individuals. The human reference genome (hg38), which does not include genetic variation or zygosity, served as a baseline sequence. For this reference setting, we paired each gene’s sequence with the average expression value across all individuals, providing a population-level baseline for comparison with individual-specific predictions.

### 2.2 Study design

#### 2.2.1 Ancestry prediction

We formulated ancestry prediction as a supervised classification problem on fixed genomic windows within the MHC region. Our goal was both to benchmark HyenaDNA embeddings for ancestry inference and to test whether explicitly modeling both haplotypes, thereby encoding zygosity, improves performance over single-copy representations.

The overall pipeline is illustrated in Figure 1a. For each individual and haplotype (A and B), a sequence of length *N* was passed independently to the HyenaDNA encoder, producing token-level embeddings of size *N* × *E*. These embeddings were reduced to a single vector per haplotype using one of three aggregation strategies: mean pooling, max pooling, or last-token embedding. This yielded one *E*-dimensional embedding per haplotype, which we then combined into different input configurations for downstream classification.

**Figure 1.**
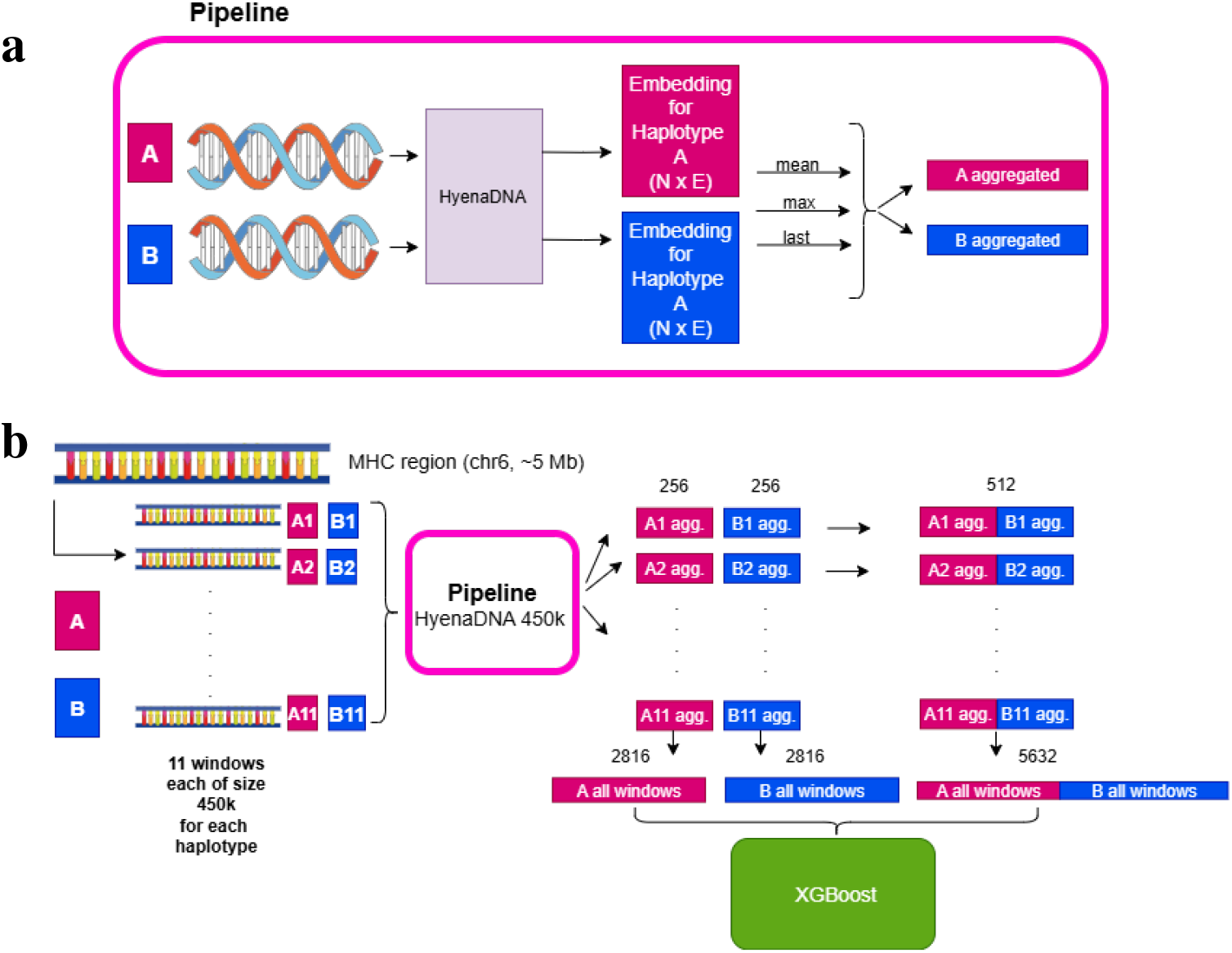
Ancestry prediction overview: **a**, pipeline for generating embeddings for haplotypes using HyenaDNA. **b**, Ancestry prediction pipeline for the MHC region showing embeddings dimension and different concatenation methods.

Mean pooling is recommended for short sequences and last-token for long sequences by HyenaDNA authors, while we also evaluated max pooling—motivated by its effectiveness in convolutional models and the hierarchical convolutional structure of the Hyena operator (Poli et al., 2023), capturing the most salient embedding features.

Figure 1b summarizes the ancestry prediction setup. The 5 Mb MHC region on chromosome 6 was divided into eleven non-overlapping windows of 450 kb. For each individual in the 1000 Genomes dataset, both haplotypes were extracted for all windows and processed with HyenaDNA using a sequence length of 450 kb. Each window produced token-level embeddings that were aggregated into a 256-dimensional vector per haplotype.

From these embeddings, we constructed three input representations:

- **Haplotype A only:** Eleven 256-dimensional embeddings (one per window) concatenated into a vector of size 2,816.
- **Haplotype B only:** Same as above, using haplotype B.
- **Concatenated AB:** For each window, the two haplotype embeddings were concatenated into a 512-dimensional vector, then concatenated across eleven windows, yielding a 5,632-dimensional representation.

These vectors were used as input features to an XGBoost classifier trained to predict the five super-populations.

#### 2.2.2 Gene Expression prediction

We formulated gene expression prediction as a regression task in which the input is a 1,000 bp DNA sequence centered on the transcription start site (TSS) and the output is the corresponding log-transformed gene expression level in LCLs. This design enables a direct comparison between models that ignore genetic variation (reference genome only) and models that incorporate individual genotypes and zygosity. We systematically varied both the encoder (CNN versus NT-based) and the sequence representation (reference only, one copy with genetic variation, two copies with genetic variation and zygosity) to disentangle their contributions.

Specifically, we extracted 1,000 bp windows centered on TSSs for all protein-coding genes. For the reference genome condition, each gene’s sequence was paired with the average expression across individuals. For the Geuvadis condition, the same sequence window was reconstructed for each individual using their diploid genotype data, and the model was trained to predict that individual’s expression level (Figure 2).

**Figure 2.**
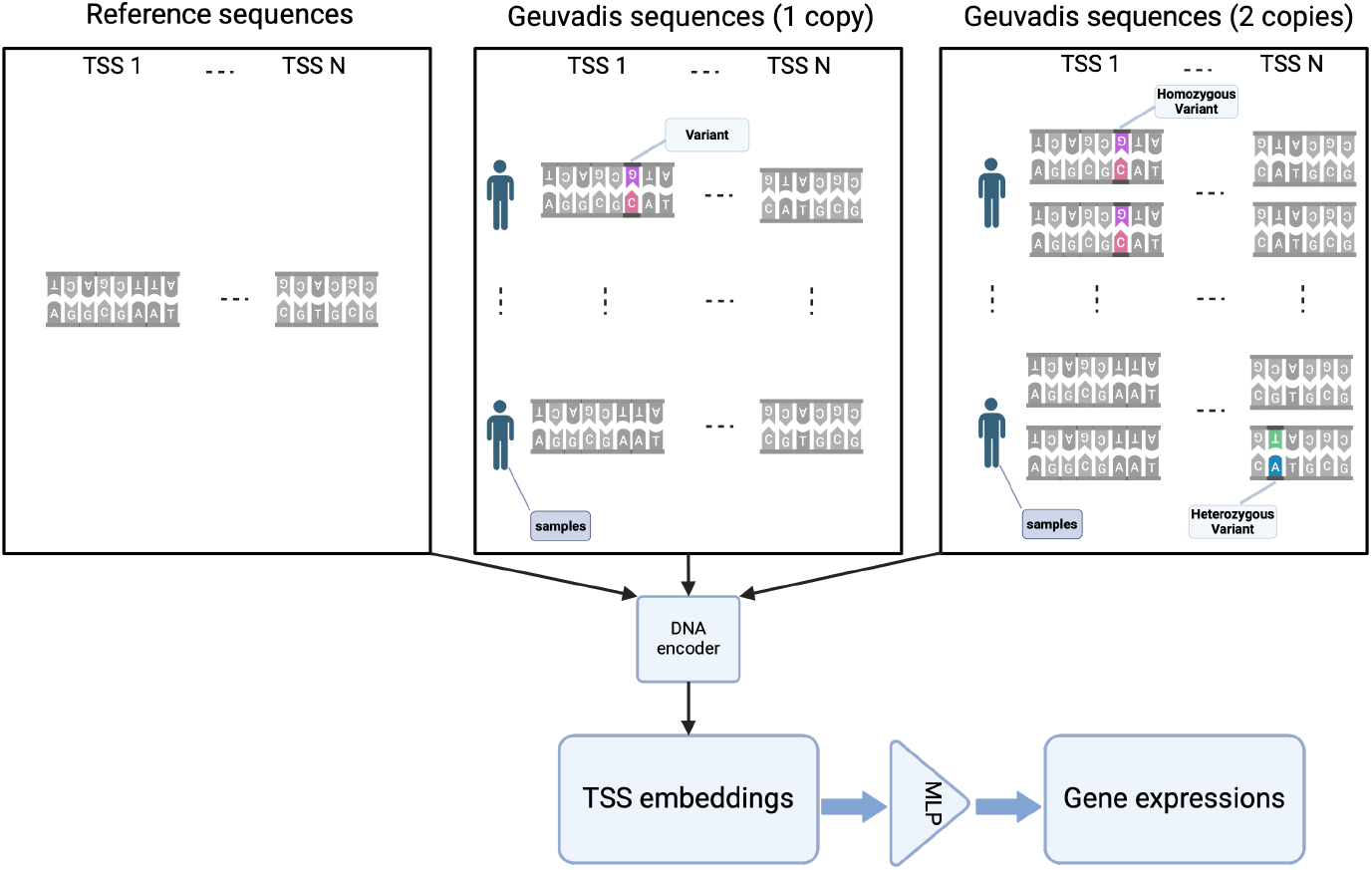
Gene expression overview: 1000 bp DNA sequences around TSS are embedded using a DNA encoder, then used to predict average (reference) or individual (Geuvadis) gene expression via a feedforward neural network.

### 2.3 DNA ENCODERS

#### 2.3.1 HyenaDNA FOR ANCESTRY PREDICTION

For ancestry prediction, HyenaDNA served as the DNA encoder. HyenaDNA introduces the Hyena operator, which achieves subquadratic time complexity 𝒪(*L* log_2_ *L*), where *L* is the sequence length, enabling long-range dependency modeling at scales of hundreds of kilobases (up to 1 Mb) without the quadratic cost of full attention. To support nucleotide-level modeling, HyenaDNA uses a vocabulary of only the four DNA bases (A, C, G, T) plus special tokens and is pretrained with a next-nucleotide prediction objective. This configuration allows HyenaDNA to represent single-nucleotide variants without relying on k-mer or BPE encodings and makes it especially suitable for modeling large diploid genomic regions with base-level resolution.

HyenaDNA also served as the architectural foundation for Evo (Nguyen et al., 2024), which extended the Hyena design to a larger scale. Because accurate ancestry prediction depends on resolving single-nucleotide variants, the combination of single-base tokenization and long-range context modeling in HyenaDNA is well matched to this task. We used the 450k-context HyenaDNA model with approximately 1.6 million parameters. Each extracted sequence was truncated or padded to a fixed length of 450,000 nucleotides. All sequences were provided as nucleotide strings (A, C, G, T), rather than one-hot encodings.

#### 2.3.2 DNA ENCODERS FOR GENE EXPRESSION PREDICTION

For gene expression prediction, we evaluated three types of DNA encoders:

- **CNN without pretraining:** A baseline model composed of three 1D convolutional layers with 4, 8, and 12 filters, respectively, a kernel size of 5, and padding of 2.
- **Nucleotide Transformer (NTREF):** A 500M-parameter model pretrained on the human reference genome.
- **Nucleotide Transformer (NT1000G):** A 500M-parameter model pretrained on a diverse set of genomes from the 1000 Genomes Project.

The Nucleotide Transformer is a DNA language model based on the BERT architecture that uses 6-mer tokenization and masked language modeling for pretraining, enabling it to capture contextual sequence patterns and dependencies. We selected this model family because it offers multiple pretrained variants, allowing a direct comparison of pretraining data (reference-only versus population-level variation) and its impact on downstream performance. Nucleotide Transformer has also demonstrated strong accuracy across a variety of genomic benchmarks in previous studies, making it a robust baseline for our experiments.

For the CNN, reference and single-copy Geuvadis sequences were represented by standard one-hot encodings. For two-copy Geuvadis sequences with zygosity, we used an additive genotype encoding in which heterozygous alleles were encoded as 1 and homozygous alternate alleles as 2 (Figure 3).

**Figure 3.**
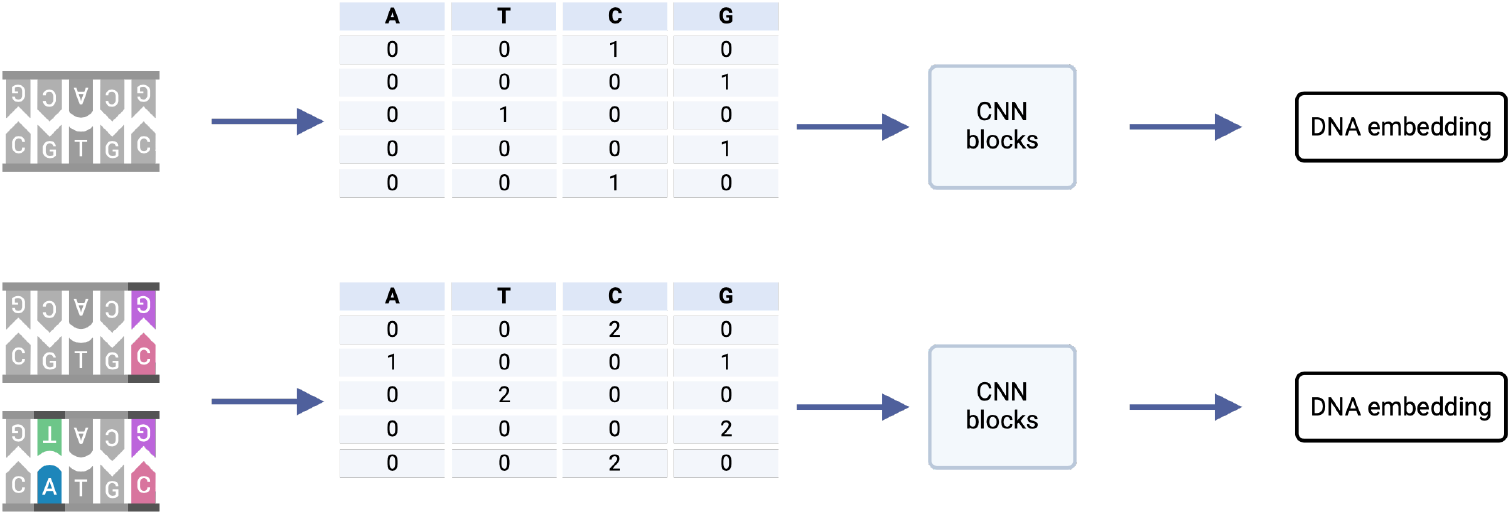
To train the CNN models, DNA sequences are first encoded using either one-hot encoding (when using reference or 1 copy of Geuvadis genomes) or additive genotype encoding (when using 2 DNA copies).

For NTREF and NT1000G, DNA sequences were first tokenized into 6-mers and passed through the respective pretrained models to generate embeddings. When using two DNA copies, we modeled zygosity by concatenating the embeddings corresponding to each copy (Figure 4).

**Figure 4.**
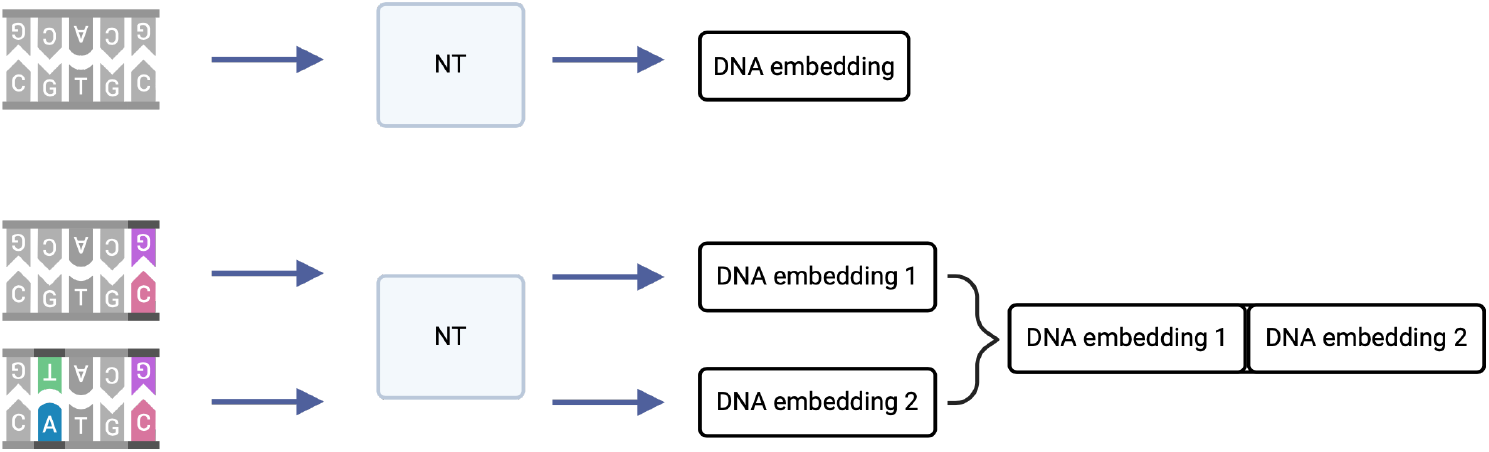
For the NT-based, we passed the DNA sequences through the respective pretrained models to generate DNA embeddings. When using 2 DNA copies, we concatenated the embeddings.

### 2.4 Model training and evaluation

#### 2.4.1 Ancestry prediction

For ancestry prediction, we trained XGBoost classifiers on HyenaDNA-derived embeddings using stratified five-fold cross-validation. Stratification preserved the original distribution of ancestry classes in each fold, reducing potential bias due to imbalanced representation of the AMR superpopulation. Within each fold, we used an 80%/20% train–validation split. XGBoost hyperparameters were chosen based on configurations previously validated for similar tasks by the HyenaDNA authors: learning rate of 0.1, maximum tree depth of 3, 1,000 estimators, and a multiclass softmax objective. We explored class weighting to further mitigate class imbalance, but this yielded negligible improvements and was not used in the final models. All classifiers were implemented using the Python XGBoost library. Performance was evaluated using macro F1-scores computed across the five superpopulations.

#### 2.4.2 Gene expression prediction

For gene expression prediction, we adopted the chromosome-based data split defined in the original workshop: chromosome 8 for testing, chromosome 9 for validation, and all remaining autosomes for training. For each encoder–representation combination, we trained a multilayer perceptron (MLP) with a single hidden layer of 64 units on top of the fixed DNA embeddings (for NTREF/NT1000G) or convolutional features (for the CNN).

Training was performed for up to 100 epochs with early stopping based on validation performance. The primary evaluation metric was the Pearson correlation coefficient between predicted and observed gene expression values on the held-out test set. Model selection and early stopping were based on validation-set Pearson correlation.

## 3 Results

### 3.1 Ancestry Prediction

#### 3.1.1 Biological relevance of MHC WINDOWS

We examined the biological relevance of the high-performing windows in the MHC region, which plays a central role in immunity and exhibits exceptional genetic diversity across populations. In Figure 5, our top-scoring ancestry prediction windows correspond to distinct MHC subregions rich in HLA and other immune genes, the details of genes and windows coordinates are provided in Appendix.

**Figure 5.**
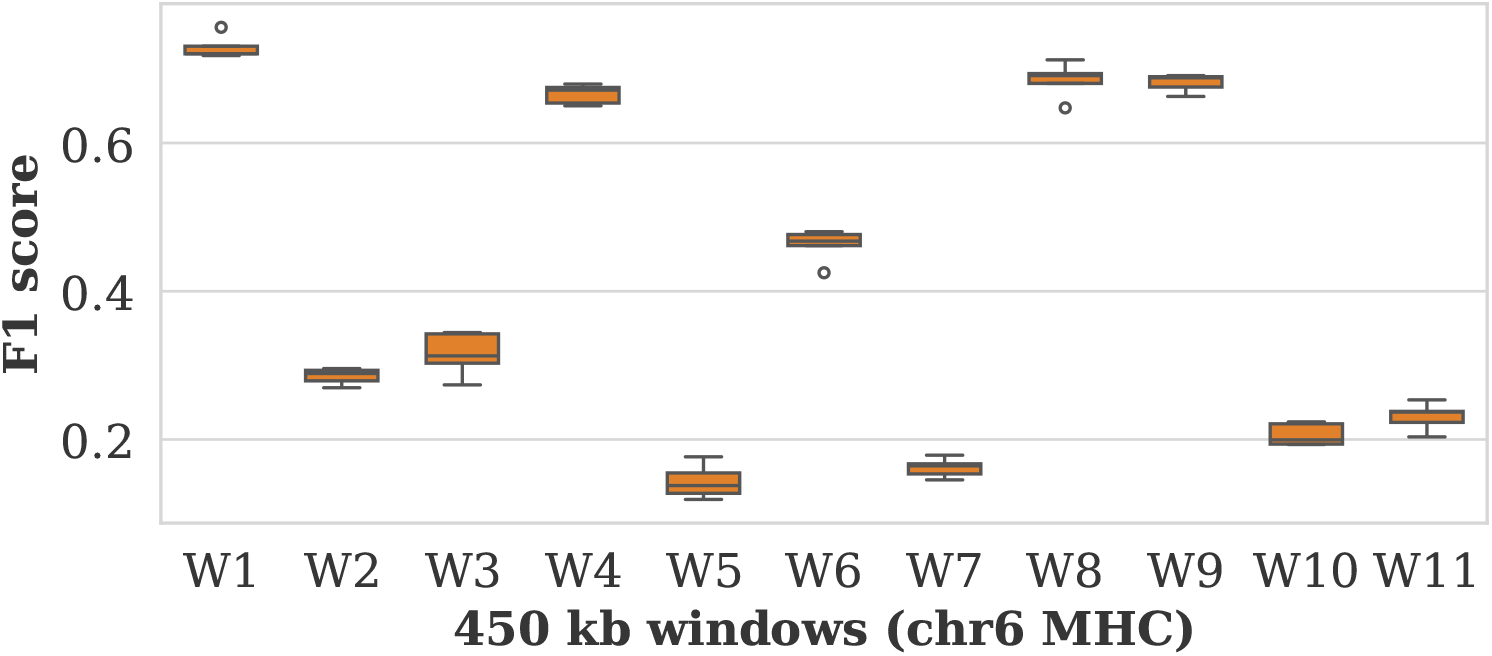
The F1 scores obtained from stratified cross-validation when applying XGBoost to the max aggregated embeddings of each 450 kb window of concatenated haplotypes embedding of size 512 in the MHC region.

This result indicate that the strongest ancestry signals arise from windows covering MHC Class I, II, and III loci, which are regions subject to intense balancing selection and highly polymorphic across global populations. Although window one exhibits the highest F1 scores, we do not find supporting literature indicating that the genes in this window vary significantly between superpopulations, thus it was not the main focus of our biological interpretation. To further characterize these regions, we quantified the number of variants in each 450 kb window and observed that window one actually contains the fewest variants overall, as shown in Figure S1a. We also counted mutations unique to each superpopulation, as shown in Figure S1b, but found no correlation between the number of these unique mutations and the classification performance of each window.

The concentration of population-specific alleles, especially within the HLA gene family and their neighboring immune genes, provides both high prediction accuracy and direct biological insight into human population structure. This agrees with previous findings that MHC variation is a primary source of ancestry differentiation (Dilthey, 2021);(Logsdon et al., 2025).

#### 3.1.2 Integrating both haplotypes improves prediction accuracy

Across all aggregation strategies (mean, max, last token), models built on concatenated haplotype embeddings significantly outperformed those using either haplotype alone. The enhancement shown in Figure 6 reflects the added information encoded by zygosity and allele-specific variation between the maternal and paternal chromosomes. On average, concatenating both haplotypes resulted in a 6.6% increase in macro F1-score over single-haplotype inputs, confirming that zygosity is a meaningful, non-redundant signal for population inference. Max pooling was the most robust aggregation strategy, likely due to its ability to preserve maximum feature variation across long windows, as smoothing via mean aggregation could hide rare but highly informative variants, and taking last token only might not capture the entire 450 kb sequence length.

**Figure 6.**
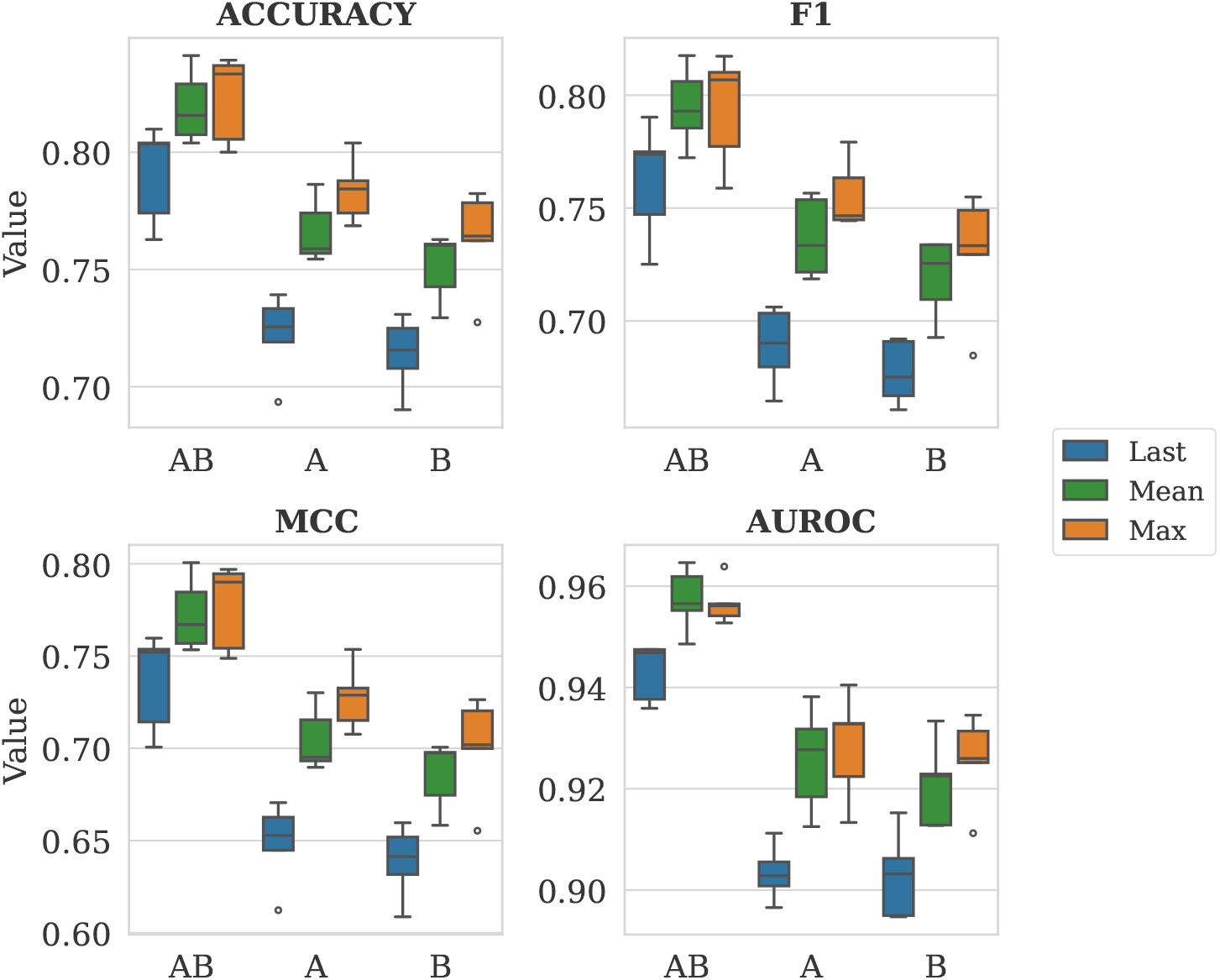
Performance of XGBoost using different aggregation strategies of HyenaDNA token embeddings (last, mean, max) and haplotype configurations (A only, B only, or A and B concatenated) for ancestry prediction. Paired t-test, F1 score comparisons: Last: *p*(AB| A) = 7.0 ×10^*−*4^, *p*(AB |B) = 3.4 ×10^*−*3^, *p*(A |B) = 0.39. Mean: *p*(AB |A) = 1.2 ×10^*−*5^, *p*(AB| B) = 6.3 ×10^*−*5^, *p*(A| B) = 0.028. Max: *p*(AB |A) = 2.0 ×10^*−*2^, *p*(AB| B) = 5.8 ×10^*−*4^, *p*(A |B) = 0.10. Embeddings were derived from 11 windows in the MHC region and evaluated using stratified 5-fold cross-validation.

#### 3.1.3 Dimensionality reduction via top windows selection

To avoid *p* >> *n* problem—that is, the scenario where the number of features vastly exceeds the number of samples (individuals)—we focus on the top four windows mentioned in the first result, enabling a substantial reduction in feature dimensionality, from 5,632 (all windows) to 2,048 (four windows). Despite the reduction, Figure 7 shows that classification performance remained high, demonstrating that the most informative population signals are concentrated within specific MHC intervals. This approach supports both interpretability and computational efficiency, prioritizing regions of strong biological relevance for ancestry prediction.

**Figure 7.**
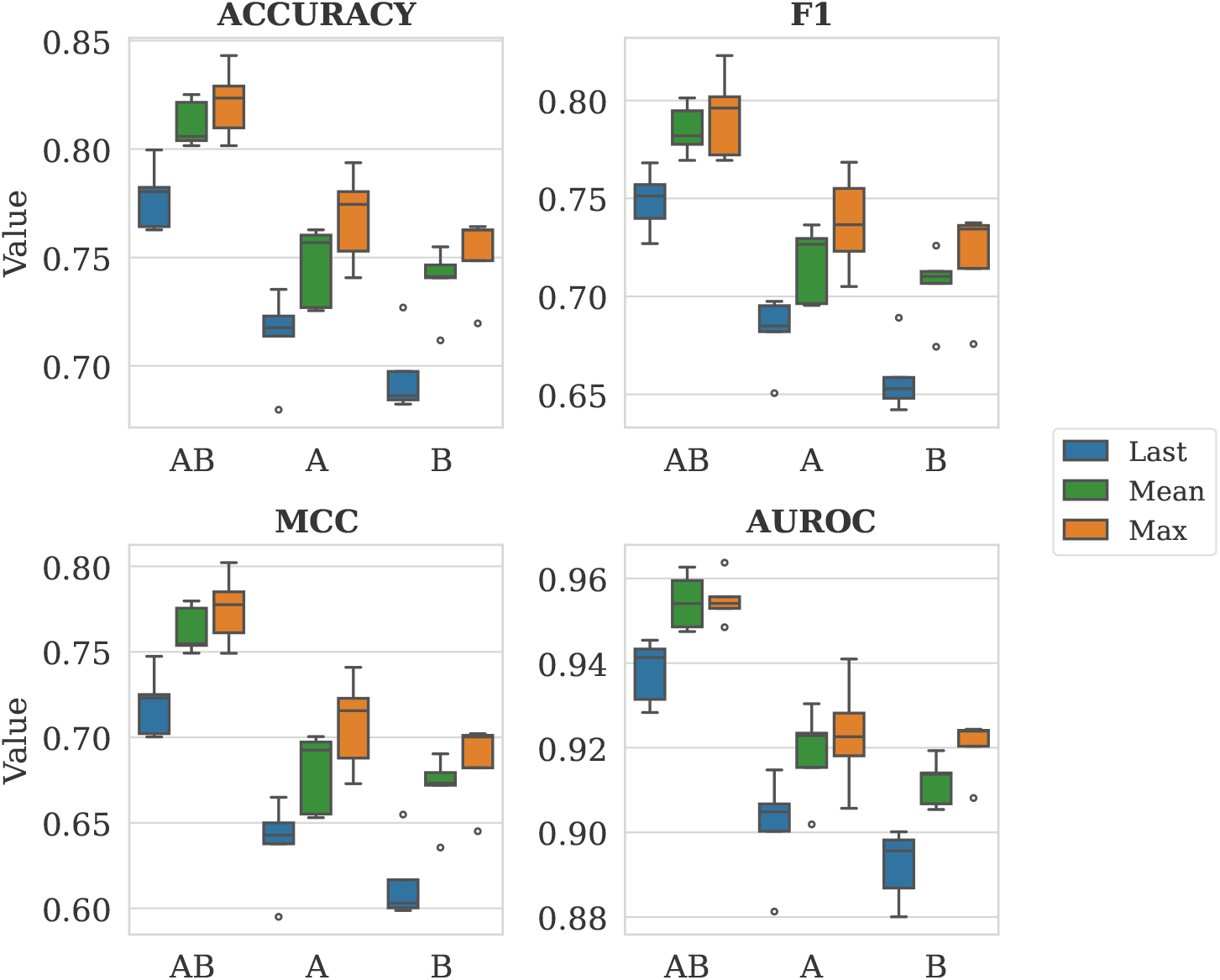
Performance of XGBoost using different aggregation strategies of HyenaDNA token embeddings (last, mean, max) and haplotype configurations (A only, B only, or A and B concatenated). Embeddings were derived from the four highest-performing windows in the MHC region; stratified 5-fold cross-validation was applied. Paired t-test, F1 score comparisons: Last: *p*(AB|A) = 2.2 × 10^*−*3^, *p*(AB|B) = 7.8 × 10^*−*5^, *p*(A|B) = 0.090. Mean: *p*(AB|A) = 8.7 × 10^*−*4^, *p*(AB|B) = 5.7 × 10^*−*4^, *p*(A|B) = 0.51. Max: *p*(AB|A) = 1.3 × 10^*−*2^, *p*(AB|B) = 2.4 × 10^*−*3^, *p*(A|B) = 0.38.

### 3.2 Gene expression prediction

We evaluated the impact of genetic variation and zygosity on gene expression prediction using CNN- and NT-based encoders. Figure 8 summarizes the Pearson correlation coefficients on the independent test set for all model and input combinations.

**Figure 8.**
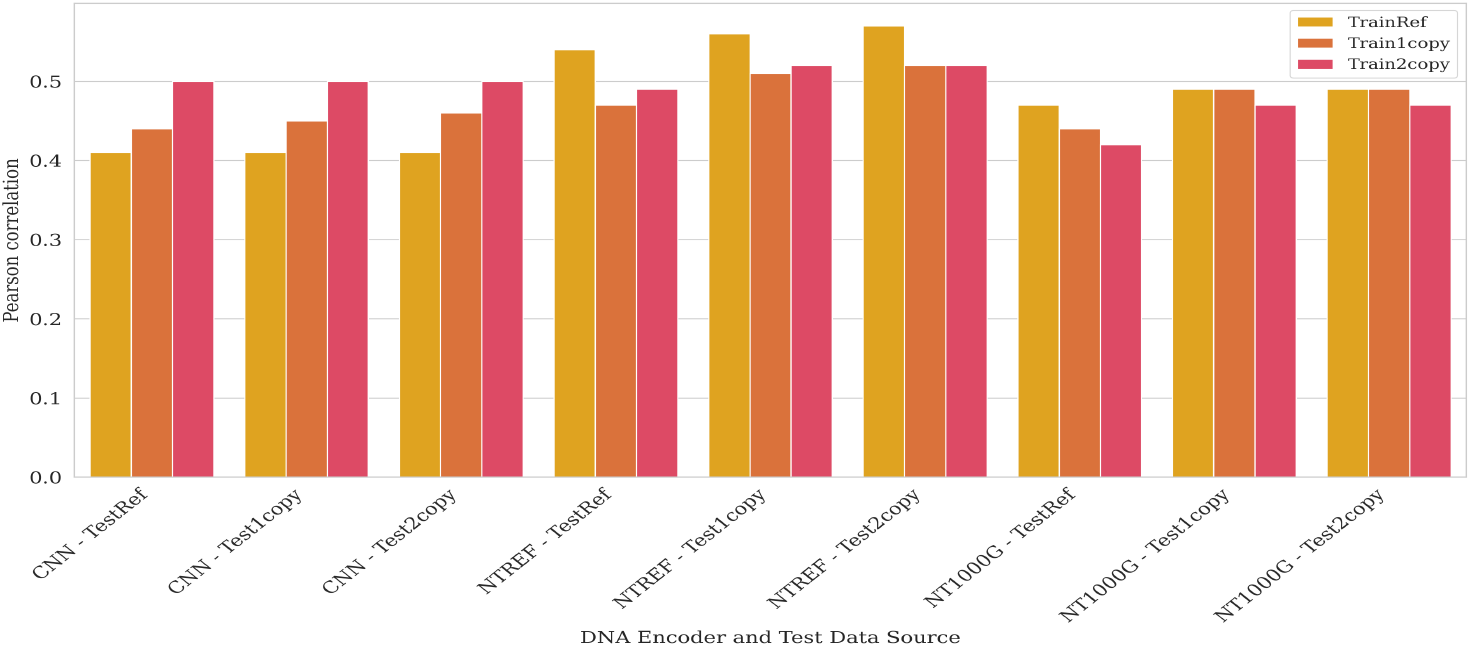
Pearson correlation coefficients of various models on the unseen test set. The X-axis represents the models and the test sequences used, while colors indicate the source of the training sequences.

For the CNN models, performance improved when training on sequences that incorporated genetic variation and zygosity compared with reference-only inputs. Using one DNA copy containing individual-specific variants increased correlation relative to the reference genome, and using two DNA copies with additive genotype encoding provided an additional gain. These results indicate that even a relatively simple architecture can exploit explicit zygosity information to capture dosage-related effects on gene expression.

In contrast, NT-based models (NTREF and NT1000G) displayed the opposite trend. When trained on inputs that included genetic variation or zygosity, their performance decreased compared with training on reference sequences only. Moreover, across all settings, NTREF consistently out-performed NT1000G, in line with previous work showing stronger downstream performance for reference-only pretraining. Together, these patterns suggest that current NT pretraining objectives, which are optimized for masked token recovery on largely reference-like sequences, may not yet fully leverage the additional complexity introduced by explicit variation-aware inputs.

## 4 Discussion

This study provides a systematic evaluation of zygosity-aware DNA representations across two complementary tasks: ancestry prediction from long-range genomic context and gene expression prediction from promoter-proximal sequences. For ancestry, we used HyenaDNA to embed phased haplotypes across the MHC region and asked whether explicitly modeling both parental copies improves classification into five superpopulations. For gene expression, we compared baseline convolutional neural networks (CNNs) to pretrained DNA language models (Nucleotide Transformer variants) on a TSS-centered prediction task, probing how different ways of encoding genetic variation and zygosity impact performance. Together, these experiments were designed to disentangle the contributions of (i) diploid versus single-copy representations, (ii) simple versus pretrained encoders, and (iii) reference-only versus variation-aware inputs for variation-sensitive downstream tasks.

Our findings show that diploid modeling captures biologically meaningful information that is missed by single-copy representations. In the ancestry prediction task, concatenating HyenaDNA embeddings from both haplotypes consistently improved macro F1-score relative to models based on either haplotype alone, across all aggregation strategies. This indicates that zygosity and allele-specific combinations across parental chromosomes encode non-redundant ancestry information, beyond what can be recovered from a single haplotype or summary variant statistics. The strongest signals were concentrated in MHC windows spanning classical HLA loci and complement genes, regions known to be highly polymorphic and subject to strong balancing selection. Focusing on these biologically enriched windows allowed us to substantially reduce feature dimensionality while preserving high classification accuracy, yielding ancestry models that are both efficient and interpretable.

In the gene expression prediction task, the way genetic variation and zygosity are encoded turned out to be as important as the choice of model architecture. Simple CNNs trained from scratch benefitted from an explicit dosage-aware encoding, where heterozygous sites are encoded as 1 and homozygous alternate sites as 2. This zygosity-aware representation improved performance over both reference-only inputs and single-copy sequences with genetic variation, showing that even relatively shallow models can exploit diploid information when it is encoded in a biologically meaningful way. By contrast, Nucleotide Transformer models did not benefit from including variation or zygosity during training, and the variant pretrained on diverse genomes (NT1000G) underperformed the reference-only model (NTREF). A plausible interpretation is that current masked language modeling objectives and reference-centric pretraining regimes bias DNA-LMs toward learning reference patterns rather than allele-specific effects.

This work has several limitations. First, the ancestry analysis is restricted to the MHC region and to only five superpopulations in the 1000 Genomes Project. While this region is a natural benchmark due to its high polymorphism and immunological relevance, it may overestimate the benefits of diploid modeling relative to more typical genomic regions with lower diversity. Second, the gene expression prediction experiments are limited to lymphoblastoid cell lines from the Geuvadis cohort and to 1,000 bp windows centered on transcription start sites. Regulatory architecture is known to involve distal enhancers and tissue-specific effects, which are not captured in our current setup. Third, our comparison of model architectures and encodings is deliberately constrained: we focus on a single CNN architecture, two Nucleotide Transformer variants, and a specific set of zygosity-aware encodings. Other architectures (e.g., transformers or state-space models trained from scratch) could lead to different relative performance.

Despite these limitations, our results point toward several promising directions for future work. At the pretraining level, there is a clear need for DNA language models that are explicitly diploid- and population-aware, for example through haplotype-specific tokenization, variant-aware masking schemes, or objectives that directly couple sequence variation to quantitative phenotypes. Recent models such as VariantFormer (Ghosal et al., 2025), which encode zygosity via IUPAC ambiguity codes, illustrate one path forward and already improve personalized gene expression prediction. Extending such ideas to long-range architectures like HyenaDNA or to multimodal settings that integrate sequence, expression, and epigenomic profiles could further enhance performance on variation-sensitive tasks. On the application side, the diploid embedding strategies explored here could be systematically evaluated for other phenotypes where zygosity is mechanistically important, such as splicing, chromatin accessibility, or molecular mechanisms of disease variants. More broadly, integrating diploid-aware representations into pipelines for ancestry inference, variant interpretation, and precision medicine may help bridge the gap between population-level genomic models and individualized predictions that respect the full diploid nature of the human genome.

## Data AND Code availability

The data supporting the findings of this study are openly available. All HyenaDNA-derived.`pkl` files used for training and evaluating the ancestry classification and gene expression prediction models are archived at Zenodo (https://zenodo.org/records/17533617), The full set of scripts for model training, evaluation, and figure generation is available in a dedicated GitHub repository at https://github.com/huRashidy/Zygosity_DNALM, where users will also find environment specifications and further usage details. All resources are versioned and permanently archived, ensuring full reproducibility of the results presented in this manuscript.

