## Supplementary figure and data for "Zygosity-Aware DNA Language Modeling Improves Ancestry and Gene Expression Prediction"

### 5 Appendix

#### 5.1 Supplementary figures

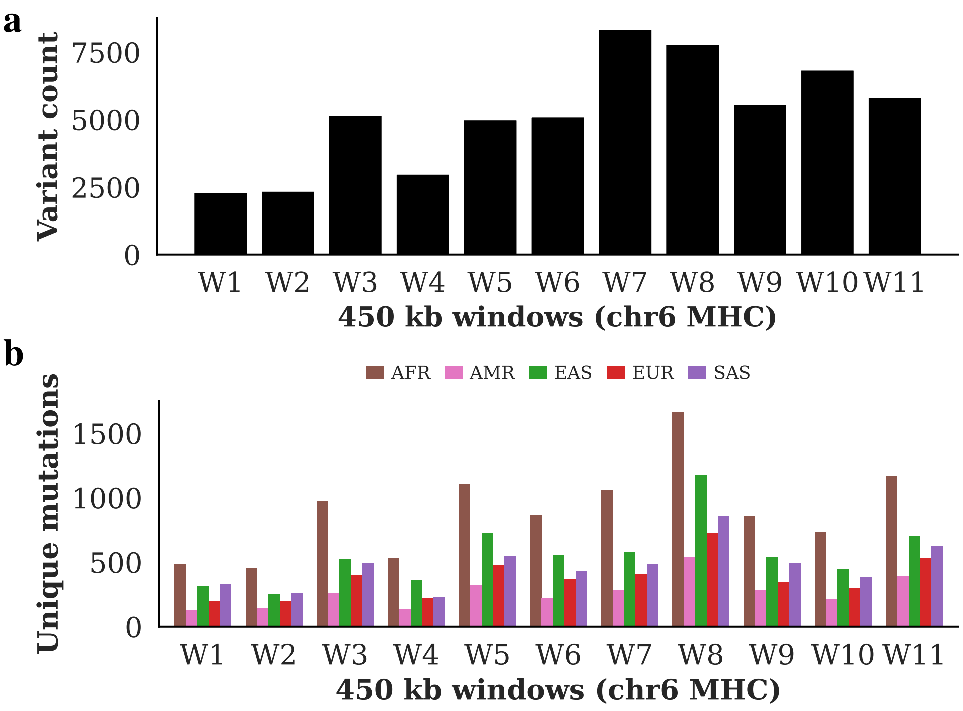


Supplementary Figure S1: **a**, The total variant counts for each 450 kb window across the MHC region (chr6). **b**, The counts of mutations unique to each superpopulation (AFR, AMR, EAS, EUR, SAS) within each window.

#### 5.2 Genes and Genomic Coordinates in Top MHC Windows

- **Window four** ($29,860,120$–$30,310,120$ bp): Contains the *HLA-A* gene, that is a key classical Class I locus, alongside additional HLA Class I genes and members of the TRIM family, which are involved in antigen processing and innate immunity.
- **Window eight** ($31,660,120$–$32,110,120$ bp): Overlaps the MHC Class III region and includes complement genes (*C2*, *C4A*, *C4B*, *CFB*), heat shock proteins (*HSPA1A*, *HSPA1B*, *HSPA1L*), and immune regulatory loci such as *STK19* and *SKIV2L*.
- **Window nine** ($32,110,120$–$32,560,120$ bp): Encompasses the *HLA-DRA* gene, a central Class II locus, as well as multiple *HLA-DRB* paralogs and immune-related genes including *NOTCH4* and *AGER*.
